# grID: A CRISPR-Cas9 guide RNA Database and Resource for Genome-Editing

**DOI:** 10.1101/097352

**Authors:** Vinod Jaskula-Ranga, Donald J. Zack

## Abstract

CRISPR-Cas9 genome-editing is a revolutionary technology that is transforming biological research. The explosive growth and advances in CRISPR research over the last few years, coupled with the potential for clinical applications and therapeutics, is heralding a new era for genome engineering. To further support this technology platform and to provide a universal CRISPR annotation system, we introduce the grID database (http://crispr.technology), an extensive compilation of gRNA properties including sequence and variations, thermodynamic parameters, off-target analyses, and alternative PAM sites, among others. To aid in the design of optimal gRNAs, the website is integrated with other prominent databases, providing a wealth of additional resources to guide users from *in silico* analysis through experimental CRISPR targeting. Here, we make available all the tools, protocols, and plasmids that are needed for successful CRISPR-based genome targeting.

## INTRODUCTION

CRISPR type II systems are naturally occurring prokaryotic immune defense mechanisms that have been repurposed for genome engineering (1–3), regulation (4–6), and other sequence-specific genomic manipulations (7). CRISPR targeting occurs through a ribonucleoprotein complex comprised of the Cas9 nuclease and the guide RNA (gRNA), a synthetic fusion between the bacterial crRNA and tracrRNA (8). The target site, formed from a spacer region followed by a Protospacer Adjacent Motif (PAM), is recognized by both components: Cas9 binds the short nucleotide PAM sequence, while the gRNA serves as the primary determinant in targeting through complementary base-pairing with the spacer region. The Cas9/gRNA complex from S. *pyogenes*, the most commonly used CRISPR system, recognizes a 20-nucleotide spacer region and the NGG PAM sequence, and therefore, the CRISPR targeting site can be represented as a contiguous sequence in the form N_20_NGG (9).

A key feature of the CRISPR RNA-directed DNA cleavage system is that Cas9 target recognition is programmable through ribonucleotide sequence changes in the guide RNA (8). While the N_20_NGG sequence forms the minimal recognition site, further constraints are generally imposed due the pol III polymerase used to synthesize the gRNA (3,10), or other sequence-dependent factors such as *T_m_*, GC content (11), or polymeric runs. Another important factor is the presence of highly similar sequences in the genome, which can lead to off-target mutagenesis. This effect, further compounded by the number and position of mismatches, can significantly affect CRISPR targeting. Thus, judicious gRNA selection remains the principal factor in successful targeting.

To aid in the design and selection of optimal gRNAs, we have created grID (<u>gR</u>NA <u>Id</u>entification), a CRISPR-Cas9 guide RNA database and resource for genome editing (http://crispr.technology/). Additionally, we have integrated within grID a mechanism for members of the research community to contribute data on their success, or lack thereof, with particular gRNAs, general comments, and suggestions for database improvements. We hope that grID will serve as an evolving database of useful resources that will aid the genome editing research community by adding to the power and variety of already existing CRISPR/Cas9 design tools (see http://omictools.com/crispr-cas9-c1268-p1.html for a partial compilation of available resources).

## MATERIALS AND METHODS

### Identification of CRISPR sites

To determine the potential set of targeting sites in a genome for a given Cas9 ortholog, engineered variant, or alternative CRISPR system, a custom regular expression Perl script was used to search for the occurrences of that sequence, allowing presence in either orientation and for overlapping sites. For example, the 23-nucleotide sequence N_20_NGG, was searched for SpCas9. Orthologous Cas9 searches were performed using N_23_GRRT for SaCas9(12), N_18_NNAGAAW for StCas9 (9), and N_24_NNNNGATT for NmCas9 (13). To search for engineered Cas9 variants with altered PAM specificity, searches were performed using N_20_NGCG for SpCas9 VRER, N_20_NGAG for SpCas9 EQR, and N_20_NGAN for SpCas9 VQR (14). The SpCas9 D1135E variant, and high-specificity/high-fidelity variants, eSpCas9 and SpCas9-HF1, are a more specific SpCas9 targeting Cas9 proteins(15,16). N_20_NAG, considered an off-target for SpCas9, was also searched. For the alternative Cpf1 CRISPR system (AsCpf1 and LbCpf1), TTTNN_23_ was searched (17). The reference genome sequences were constructed by concatenation into a single chromosome file obtained from the repeat-masked assembly sequence; sequences masked by N’s were purposely excluded by the Perl regular expression, and thus excluded from the initial set of CRISPR sites. The reference genomes used were: hg19 for Human (NCBI build GRCh37), mm10 for Mouse (NCBI build GRCm38); rn5 for Rat (RGSC Rnor_5.0), danRer7 for Zebrafish (Sanger Institute Zv9), ce10 for C. *elegans* (WormBase v. WS220), and sacCer3 for S. *cerevisiae* (SGD). Subsequent text manipulations were performed using standard Unix, AWK, and/or Perl scripting. grID numbering is assigned incrementally starting with the first chromosome. The total set of CRISPR sites is available for download as either FASTA formatted or BED formatted files (http://crispr.technology/data/).

### Thermodynamic Properties

∆H° and ∆S° parameters for nearest neighbor DNA/RNA hybrid formation (18) were used to calculate the *T*_m_ from the free energy derived equation: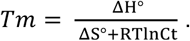 RNA secondary structure analysis of the spacer was performed using RNAFold (19) after converting the protospacer sequence to ribonucleotides, the dot-bracket notation and ∆G values were databased.

### Off-target analysis

Potential genomic off-target sites were predicted using Bowtie (20), allowing up to three mismatches along the target site (-v = 3), while suppressing the output for any sites reporting more than twenty-five alignments in the genome (-m = 25).

### Paired Nickase/Offset nicking

The offset is defined as the distance between the internal ends of the CRISPR targeting site, while the 5’ overhang length is calculated as the distance between the two nick sites (21).

### Data sources

Annotations for genomic, transcript, and protein sequences, as well as gene descriptions, were obtained from the NCBI RefSeq Database (http://www.ncbi.nlm.nih.gov/refseq/) (22), and through the UCSC genome browser (23). Sequences and annotations are locally databased and accessed using BioPerl (24). Restriction enzyme analysis is performed through Perl scripting using the target sites from REBASE (http://rebase.neb.com/rebase/rebase.html) (25). OMIM and disease data was obtained from OMIM (Online Mendelian Inheritance in Man, OMIM. McKusick-Nathans Institute of Genetic Medicine, Johns Hopkins University (Baltimore, MD, USA), 2013. World Wide Web URL: http://omim.org/) (26). dbSNP human Build 142 based on GRCh37/hg19 was used as the data source for SNPs and variation (27).

### UCSC genome visualization

The grID data in the BED file format was converted to compressed binary indexed BigBed file using the bedToBigBed Kent utility (28). Visualization through the UCSC genome browser (http://genome.ucsc.edu).

### Construction and implementation of the grID database

The grID data is stored in a MySQL relational database installed on a Unix-Apache web server. Server-side scripting is performed using Perl, while dynamic and client-side scripting is performed using JavaScript and jQuery. The front-end HTML, CSS, and JavaScript interface was developed on the Twitter Bootstrap framework and is accessible through desktop, tablet, and mobile devices.

### Plasmid construction

Plasmids encoding the H1 promoter::*Avr*II::gRNA scaffold or U6 promoter::*AvrII*::gRNA scaffold were constructed by gBlock synthesis (IDT) and Gibson assembly (NEB) (29) into pUC19 digested with *NdeI* and *XbaI* restriction enzymes. For *in vitro* gRNA expression, T7, T3, or SP6 bacteriophage promoters were similarly constructed by Gibson assembly using gBlock synthesis cloned into pUC19 (*NdeI*/*XbaI*). The promoter::target::gRNA scaffold unit is flanked by an upstream 5’ *ClaI* site and a 3’ *NotI* site; the latter is used to linearize the vector generating a 5’ overhang.

## RESULTS AND DISCUSSION

### grID system

In order to provide a universal CRISPR naming system, we introduce grID (**gR**NA **Id**entification), a nomenclature system based on the systematic identification of CRISPR-Cas9 sites. CRISPR recognition sites in the genome are uniquely designated by a two-letter prefix to indicate the species (e.g. hs for *Homo sapiens*) followed by a sequentially assigned numerical digit (e.g. <u>hs025713228</u>). The grID system provides a simple and unambiguous framework for the identification of CRISPR sites in any genome. Furthermore, the identification system is flexible enough to accommodate PAM sequences from other existing orthologs (1,12,13), engineered or evolved Cas9 variants (14–16), and alternative CRISPR systems (17) (**Supplemental Table 1**).

We began by determining the potential CRISPR sites in the human genome for the S. *pyogenes* Cas9 by searching for all occurrences of the sequence N_20_NGG. Since for most genome-engineering applications targeting within repetitive regions is undesirable, we suppressed such loci by using the repeat-masked sequence for the initial genome reference. In total, there are 137,409,562 potential CRISPR sites that fit these criteria within the human genome (<u>hs000000001</u> - <u>hs137409562</u>), and their location and sequence are easily accessible through the grID database.

Off-target analysis is a major component of the database, comprising the computationally predicted off-targets for grID entries. The off-target parameters are pre-calculated and databased to avoid the computational time that users would otherwise expend while performing analysis; however, to balance virtually instantaneous data retrieval with database size, we limited the pre-calculated analysis to 0, 1, 2, or 3 mismatches in CRISPR sites totaling less than 25 predicted off-target loci. Sites with 25 or greater off-target loci are still indicated; however, users are generally advised against using such sites. For example, in the human genome we first performed a genome-wide alignment for the 137,409,562 sequences in the grID database allowing for up to 3 mismatches along the entire targeting sequence, which resulted in 10,172,371,379 potential loci. By limiting the results to sequences with 25 or less loci, the total number was reduced to 1,575,631,967 off-target sites. We find that this provides ample data for the vast majority of researchers, as users are unlikely to perform sequencing analysis and verification for a large number of loci. Additionally, those who are highly concerned with off-targets are likely to use more discriminating variants like the D1135E SpCas9 variant (14), or the high-specificity/high-fidelity variants (eSpCas9 and SpCas9-HF1) that exhibit low-to-undetectable off-target mutagenesis (15,16).

By querying the database against a small set of experimentally determined CRISPR sites (30), we found that the database performs accurately on all sites (**Supplemental Table 1**). Of the 6 total sites that were examined, 4 were reported to elicit a high-level of off-target mutagenesis; the two sites that were not reported to have off-target hits (<u>hs008507971</u> and <u>hs019839404</u>) also do not produce any user warnings from the database. 2 of the remaining 4 sites (VEGFA T2 and VEGFA T3) are suppressed because they intersect with repeat-masked regions, and desirably, these potentially confounding sites are absent from user queries. The remaining 2 sites produce user warnings for potential off-targets due to the presence of polymeric runs, dinucleotide repeats, and high GC content (<u>hs106331938</u>), or due to the presence of a trinucleotide repeat (<u>hs064758550</u>).

As dissociation kinetics governs Cas9 cleavage (31), we reasoned that relative target site uniqueness toward the high-affinity PAM proximal region could be used to modulate on- and off-rates. The grID Specificity is based on an approach we developed to classify gRNAs according to their relative uniqueness in the genome through iterative 5’ truncations and genomic alignments. For example, with SpCas9 the target sequences (N_20_NGG) were first aligned to the genome to identify unique sites. Next, the 5’ nucleotide was removed and the sequence (now N_19_NGG) was realigned. Those that failed to align uniquely were scored and removed from the next cycle. This process identifies more unique CRISPR targeting sites in the genome with a bias towards the seed region, which we score as “low,” “medium,” or “high” specificity. We also include other on-target scoring algorithms pre-computed for each gRNA, where applicable (32,33).

## DATABASE

The grID database provides several query options. In addition to searching by grID (e.g. <u>hs025713228</u>), users can search by NCBI reference sequence (RefSeq) ID (e.g. <u>NM 002046</u>), Gene Symbol (e.g. <u>GAPDH</u>), or any valid 23-bp targeting sequence in the form N_20_NGG (e.g. <u>GCATCTTCTTTTGCGTCGCCAGG</u>). User queries from Gene Symbols with multiple splice variants are directed to an isoform page that provides links to a unique RefSeq ID.

### grID Page

Every grID page provides an extensive set of properties that are useful in gRNA selection and as reference information:

i. A Summary section provides an overview that contains the CRISPR target sequence, grID specificity (low, medium, high), and Cas9 species of origin (e.g S. *pyogenes*) (**Figure 1A**).
ii. The Genomic Context indicates the exact genomic coordinate (chr:start-end) and strand (+/−) orientation for the target site. Additionally, the location is mapped against RefSeq coordinates, providing a descriptive location (exonic, intronic, or intergenic) (**Figure 1B**). The nearest genes subsection indicates the genes closest to the target site.
iii. The Browser section provides a simple graphical and interactive visualization tool. The embedded Jbrowse (34) frame found on each page automatically loads to the specific region of interest (**Figure 1C**).
iv. The Sequence Analysis section details the transcribed RNA sequence (gRNA spacer sequence), uniqueness status of the target (protospacer) sequence (unique or non-unique), the PAM motif, and the presence of either homopolymeric runs or pol III terminator sequences (yes or no) (**Figure 1D**).
v. A Thermodynamic Properties section includes GC-content, melting temperature (*T*_m_), and RNAfold prediction and stability. These properties are qualified (e.g. “very low,” “low,” “moderate,” “high,” and “very high”) to provide the users with some guidance for potentially problematic targeting sequences (**Figure 1E**).
vi. Users are provided with a general summary of the mismatches and numbers of off-target sites, along with detailed alignments, and mismatch location: distal (position 20-16), distal seed (position 15-11), proximal seed (position 10-1), and PAM N mismatches (**Figure 1F**).
vii. A Paired Nickase/offset nicking section is available for users who prefer to use the Cas9 (D10A) nickase (5,21). The potential gRNA mates compatible with the Cas9 (D10A) mutant, along with the respective offset distance and 5’ overhang length, are ranked by increasing distance.
viii. A Sequence section provides users with quick access to the flanking genomic sequence. 500bp of the reference sequence upstream and downstream of the target site is shown (**Supplemental Figure 1A**).
ix. The Screening Analysis section, which indicates the presence of restriction sites that fall within the targeting region, is available to aid in downstream CRISPR experimental analysis. Restriction sites, particularly those that overlap the cutsite, can be used as a convenient alternative to mismatch detection and quantification by the standard Surveyor or T7 endo I assay (35,36) (**Supplemental Figure 1B**).
x. The SNPs and Variations section provides a quick overview of dbSNP142 data within the target site that could lead to potential sequence differences (27). Minor sequence variations can explain recalcitrant targeting or be exploited for allele-specific targeting (**Supplemental Figure 1C**).
xi. The Notes section provides user alerts about gRNAs that are potentially problematic. Warnings for properties that can contribute towards a higher propensity of off-target mutagenesis are noted, such as non-unique targeting sequences, homopolymeric runs, high *T*_m_, or high GC content. Additionally, potential downstream pitfalls like strong secondary structure within the spacer or the presence of a pol III terminator are also indicated **(Supplemental Figure 1D)**.
xii. A Reference section is provided on each grID Page, allowing for the cataloging of citations publishing a target sequence. A deposit form will be available on the website, encouraging authors or general users to contribute references to the database. This in turn provides a valuable resource for the community, as well as a potential mechanism for author recognition through citations.
xiii. Finally, in an effort to promote scientific collaboration and to disseminate valuable information to the CRISPR community, we encourage use of a Comments section that is provided at the bottom of each grID page. Comments made on a specific site are only available through that grID Page, ensuring that only pertinent information is displayed to the user.

**Figure 1.**
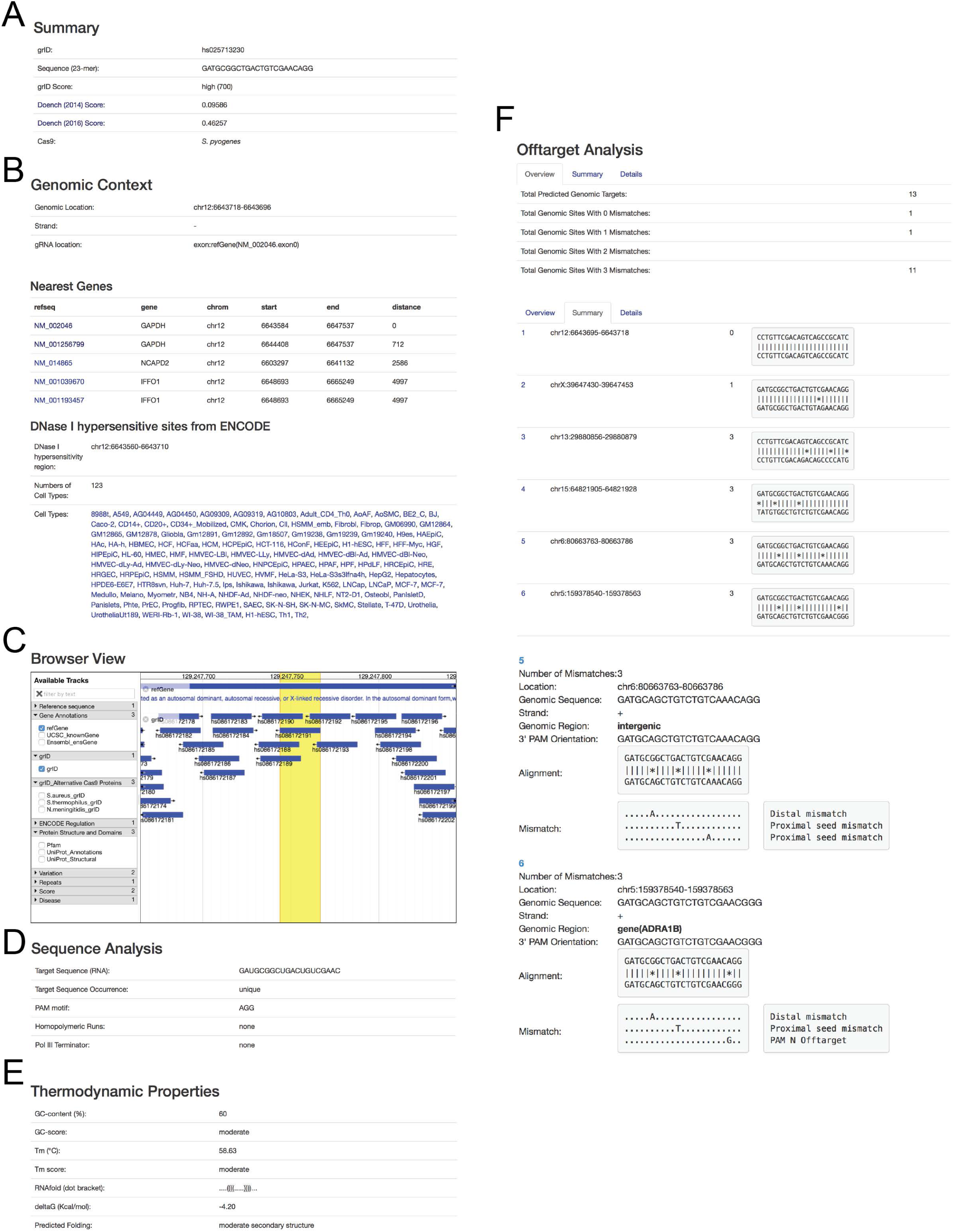
Overview of the grID page. A summary of the CRISPR target sequence and on-target score from various algorithms (**A**), genomic context of the target site (**B**), an embedded genome browser (**C**), sequence analysis (**D**), thermodynamic properties (**E**), and off-target analysis (**F**).

### Other Resources, Plasmids and Protocols

gRNA selection is only one step in the process of genome editing. Since an important aspect of the grID database/website is to create an encompassing CRISPR ecosystem available through a single centralized location, and to facilitate CRISPR research by bridging *in silico* analysis with laboratory resources, we provide additional resources to help potential users.

Similar to the grID page, the website provides useful information from the context of the gene: (i) The Summary section provides a brief gene description and summary from RefSeq (22), as well as quick links to other databases; (ii) a Disease Information section with information from the OMIM database is provided (26); (iii) a Browser section that indicates a similar graphical and interactive visualization tool as the grID page; (iv) a Genomic Context section for genome coordinates; (v) Sequence information to provide quick access to genomic and protein sequences; (vi) an interactive search and filter to identify high scoring gRNAs targeting specific exons; (vii) Notes, which provide useful additional gRNA-specific information, (viii) References, and (ix) a Comments section analogous to that described for the grID page.

The advanced search page (http://crispr.technology/search.html) allows users to target specific regions in the genome that would be useful for a wide range of CRISPR experiments. For example, users can target exons, introns, 5’ or 3’ UTRs and promoter regions, and application-specific searches, like CRISPRi (37,38), are available as well. For users who prefer data visualization through a genome browser, grID data can be viewed through a full-page JBrowse genome browser (http://crispr.technology/browser/)(34), or via the UCSC genome browser (http://crispr.technology/genomes.html)(23). Both options allow users to interact with their own data. The database and resources are primarily geared towards human and mouse targeting, however other model genomes such as rat, zebrafish, *C. elegans,* and S. *cerevisiae* are also available.

Though our resource page users have access to extensively detailed protocols that we have tested and optimized for generating specific CRISPR targeting constructs. We generated several plasmids using common *in vivo* promoters (H1 and U6) and *in vitro* promoters (T7, T3 and SP6) to facilitate gRNA transcription, and all of these are freely available through Addgene (www.addgene.org) (**Supplemental Sequences**) (10). An oligonucleotide builder (http://crispr.technology/resources/oligoform.html) is available to generate primers for cloning any gRNA in the database into various expression vectors. Construct generation for the user is simplified though the use of a common restriction enzyme (AvrII), cloning method (Gibson Assembly) (29), antibiotic selection marker (ampicillin), and sequencing primers (M13R). Optimized experimental protocols including for cell line transfection (http://crispr.technology/resources/transfection.html) and quantification of genome-editing efficiency (http://crispr.technology/resources/quantification.html), or tools useful for common genome-editing calculations(http://crispr.technology/resources/calculations.html) are also available.

Our goal is to maintain grID as a dynamic database serving the genome editing community, and efforts will be made to expand and modify the database as needed to keep pace with the rapid evolution of CRISPR-Cas9 technology.

